# Triploid asexual freshwater snails grow faster than sexual diploid conspecifics regardless of dietary phosphorus availability

**DOI:** 10.64898/2026.06.19.733397

**Authors:** Briante Najev, Zachary Minthorn, Seth Gordon, Joshua Bliss, Chase McInville, Vinny Chloros, Waad Abdella, Maurine Neiman, Amy C. Krist

## Abstract

The number of chromosome sets per nucleus is a fundamental trait, but why this number is nearly always two for multicellular eukaryotes remains unclear. Chromosomes are made of nucleic acids, which possess abundant phosphorus (P). Therefore, producing new chromosomes, as well as generating new cells and organismal growth, demands substantial phosphorus. Yet, because P is often limiting in nature, P availability could influence the prevalence of diploidy versus polyploidy. Here, we compare growth rates of diploid and triploid *Potamopyrgus antipodarum*, a freshwater snail, relative to P availability. Because diploid *P. antipodarum* are obligately sexual while obligately asexual individuals are polyploid, costs associated with sensitivity to P limitation in polyploids could also help explain the maintenance of sexual *P. antipodarum*. We raised juvenile diploid and triploid snails on either P-adequate or P-deficient diets and found that independent of P availability, juvenile triploid asexual snails grew faster and harbored higher P content as adults than sexual diploid conspecifics. Together, these results suggest life-history advantages of polyploidy or asexual reproduction that exacerbate rather than ameliorate the cost of sex. These outcomes suggest that P availability is unlikely to be a main driver of ploidy polymorphism or the maintenance of sex in *P. antipodarum*.

## Introduction

The overwhelming predominance of diploidy among eukaryotic taxa, despite frequent ploidy elevation, suggests major benefits associated with relatively low ploidy level (Wendel 2015, Wang et al. 2021). Why diploidy confers such profound advantages is not well understood, but because direct and indirect consequences of ploidy elevation affect virtually all levels of biological organization, ploidy should influence many aspects of organismal phenotype (Morris et al. 2024). Indeed, diploids have been shown to possess advantages, in at least some conditions, relative to polyploid conspecifics ranging from frogs (Chen et al. 2025) to zebrafish (van de Pol et al. 2021) and angiosperms (Guignard et al. 2016) to yeast (Gerstein et al. 2017). However, polyploids can also possess advantages over diploids. For example, triploids grow faster than diploid counterparts across multiple bivalve taxa (reviewed in Piferrer et al. 2009), and triploid flatworms (*Schmidtea polychroa;* D’Souza et al. 2005) have higher fecundity than diploid conspecifics. These patterns of polyploid advantages also extend to plants, with examples from grasses (Godfree et al. 2017), duckweed (Turcotte et al. 2024), and flowering plants (Münzbergová and Skuhrovec 2017). Here, we address the possible advantages and disadvantages of polyploidy relative to nutrient availability.

Multiple studies across a diverse range of taxa indicate that polyploids and other taxa with relatively high nuclear DNA content (e.g., via relatively large haploid genome size) may need more resources, including key limiting nutrients, than diploids to successfully meet life-history milestones. For example, primary productivity of tetraploid *Heuchera cylindrica* (poker alumroot) is more sensitive to limited nitrogen and phosphorus availability than diploid conspecifics (Anneberg and Segraves 2023). *Candida albicans* (yeast) maintains a smaller genome size when grown in a phosphorus (P)-depleted medium than when grown in media with adequate P (Gerstein et al. 2017). Similarly, the cyanobacterium *Synechococcus elongatus* loses genome copies across generations when phosphate is depleted but maintains more genome copies when the medium is P rich (Riaz et al. 2022).

Polyploids may also be more likely to experience P limitation than diploids because: 1) polyploid cells possess more DNA than diploid cells and DNA is ∼9% phosphorus (P) (Elser et al. 1996), 2) polyploid organisms can possess more phosphorus per unit mass than their diploid conspecifics (e.g., Neiman et al. 2009) and 3) therefore polyploids are more likely to require more P than diploid conspecifics. Additionally, because P is frequently limiting in nature (Cross et al. 2005, Elser et al. 2007, Du et al. 2020), freshwater organisms frequently face P limitation. While higher sensitivity to P limitation in polyploids, relative to diploids, has been reported in a wide range of eukaryote taxa (e.g., plants: Veselý et al. 2013, Anneberg et al. (2020, 2023); animals: Neiman et al.2013b, Jeyasingh et al.2015, Gerstein et al. 2017), we still know relatively little about nutrient-ploidy connections in animals relative to plants (e.g., Neiman et al. 2017).

We also consider the potential for benefits associated with polyploidy in animals when P availability is sufficient. Multiple lines of empirical evidence are consistent with this possibility. Relevant studies demonstrate that polyploid *Daphnia pulex* (water flea) grow more rapidly in P-adequate environments than diploids (Jeyasingh et al. 2015), show that tissue regeneration is more rapid in polyploid vs. diploid snails (Krois et al. 2013) and salamanders (Saccucci et al. 2016), and reveal higher growth rate and earlier reproductive maturity in polyploid vs. diploid *Potamopyrgus antipodarum* fed a P-adequate diet (Larkin et al. 2016, Gibson et al. 2017). Also, endopolyploid tissues possess higher mRNA, protein production, and growth rates than diploid tissues across both animals and plants (reviewed in Neiman et al. 2017, Morris et al. 2024) further suggesting polyploid advantages, at least in some contexts. Together, these studies suggest the potential for fitness advantages of triploids relative to diploids in P-adequate conditions.

We explored possible disadvantages and advantages of polyploidy related to P availability in diploid sexual and triploid asexual *Potamopyrgus antipodarum,* a New Zealand freshwater snail. Native populations of *P. antipodarum* are characterized by frequent coexistence among obligately sexual diploid and obligately asexual polyploids (i.e., triploid and tetraploid) along with temporally stable ploidy-level variation across populations (Lively and Jokela 2002, Neiman et al. 2011, Paczesniak et al. 2013). A survey of New Zealand lakes demonstrated wide across-lake variation in the P content of periphyton, the primary diet of *P. antipodarum* (Krist et al. 2016). Krist et al. (2016)’s survey also revealed that in most of these lakes, P was likely to be limiting for *P. antipodarum* because the periphyton C:P was above its threshold elemental ratio (TER) of 270 (defined for one triploid lineage of *P. antipodarum;* Tibbets et al. 2010). At the TER, life-history traits switch from being limited by one element to another element (Frost et al. 2006). Accordingly, because C:P in the periphyton of New Zealand lakes inhabited by *P. antipodarum* is often higher than its TER suggests that P limitation could differentially shape the expression of life-history traits like growth rate in natural populations of *P. antipodarum*.

If polyploids are more likely to experience P limitation than diploids (Lewis 1985, Cavalier-Smith 2005, Guignard et al. 2017, Neiman et al. 2017; reviewed in Hessen et al. 2010), then growth rate of triploids should be more reduced under P limitation than in diploids. However, if polyploids possess growth rate-related advantages when P is adequate, then we might expect more rapid growth from triploids in P-adequate environments than diploids. Because asexual animals and plants are often polyploid (Otto and Whitton 2000, Neiman and Schwander 2011), our focus on diploid sexuals vs. triploid asexuals also allows us to address whether potential nutrient-related disadvantages and advantages of polyploidy could help to explain the maintenance of sexual reproduction (Neiman et al. 2013a).

We addressed these knowledge gaps by experimentally manipulating dietary P availability to assess the effects of P limitation and ploidy level on body P content and individual growth rate in diploid sexual and triploid asexual *P. antipodarum*. Specifically, we asked whether the P content of snails differed by ploidy level or dietary-P treatment; triploids should only possess greater sensitivity to limited P availability if they have higher P content than diploids. To examine the potential disadvantage and advantages of polyploidy, we also addressed whether 1) growth of triploids are also more severely affected by P limitation than diploids, 2) triploids consistently grow faster than diploids when P is not limiting, and 3), if we find more rapid growth in triploids, whether this growth-rate advantage of asexual polyploid *P. antipodarum* disappears when P is limiting. These last two objectives address whether potential advantages of polyploidy, like rapid growth, depend on dietary P availability, with consequences for competitive outcomes between snails with different ploidy levels (and reproductive modes).

## Material and Methods

### Overview

Previous studies have demonstrated objective distinctions between groups of *P. antipodarum* (i.e, ploidy level: Neiman et al. 2013b; invasive vs. native status: Neiman and Krist 2016; source population: Krist et al. 2014, Krist et al. 2017) that drive different responses to P limitation but also revealed extensive variation associated with genetic background. Because our current study addresses distinctions between sexual diploid vs. asexual triploid *P. antipodarum*, we designed this experiment to control, as much as possible, for these “background” genetic effects. Thus, we focus on comparisons between two similarly diverse sets of diploid sexual and triploid asexual snails to accommodate this concern and allow us to exclude higher genetic diversity, per se, as an explanation for potential differences between diploid sexual vs. triploid asexual individuals.

Importantly, we acknowledge that we cannot decouple effects of ploidy from reproductive mode in our experiment because diploid *P. antipodarum* exclusively reproduce sexually and triploid (and tetraploid) *P. antipodarum* exclusively reproduce asexually (Phillips and Lambert 1989, Wallace 1992, Neiman et al. 2011). Nevertheless, because our experiment addresses costs of polyploidy that are not obviously directly connected to reproductive mode, and considering previous evidence for heightened sensitivity to limited P in tetraploid vs. triploid *P. antipodarum* (Neiman et al. 2013b), we hereafter focus on “ploidy”, while recognizing that ploidy is inseparable from reproductive mode in this system. However, because ploidy and reproductive mode are confounded in our system, like many other mixed sexual/asexual systems (e.g., Otto and Whitton 2000), our study is potentially relevant to understanding the maintenance of sex.

Below we describe how manipulated dietary P content of diploid and triploid snails to assess the effects of ploidy and P availability on individual growth rate and how we destructively sampled snails to measure their P content at the end of the experiment. In contrast, in the Results and Discussion we address snail body P content *before* the diet experiment because only if triploids possess higher P content than diploids would we expect greater sensitivity of triploids to limited dietary P availability.

### Snail Selection

To obtain the diploid sexual snails, we selected juveniles from a laboratory population descended from a diverse mix of ∼50 sexual male and female *P. antipodarum* collected from lakes around the South Island of New Zealand, with additional sexual individuals added to the culture every year or two from field collections. We used this “outbred sexual” population so that we could reasonably consider each individual to represent a genetically distinct replicate of a diploid sexual *P. antipodarum.* We compared these diploid snails to juvenile snails from 36 distinct and independently laboratory-cultured triploid lineages founded by individual asexual triploid females collected from lakes around New Zealand. Prior to the experiment, all snails were cultured in plastic tanks filled with carbon-filtered tap water in a 16°C room with a 16:8-hr light:dark cycle and fed *ad libitum* with powdered Sera Micro Nature ultra-fine fry fish food.

At the start of the experiment all snails were juveniles (< 2.35 mm, Jokela and Lively 1995); we measured shell length of each snail from shell aperture to apex under a dissecting microscope (Nikon SMZ800) with digital calipers. We measured P content and growth of female snails only because triploids are nearly all female (Neiman et al. 2012). Therefore, at the start of the experiment, we included approximately twice as many diploid individuals as triploid individuals because juvenile *P. antipodarum* cannot be reliably sexed (e.g., Jokela and Lively 1995), and assuming that approximately half of the diploid snails were males. At the end of the experiment, the diploid sexual snails were large enough to sex (i.e., > 3.0 mm; e.g., Larkin et al. 2016), allowing us to exclude males from data analysis.

### Diet manipulation experiment

For eight weeks, we fed half of the snails (N = 93) a low-P diet (∼458 C:P) and fed the other half (N = 93) a high-P diet (∼137 C:P). Diets are defined as limiting (low P) or sufficient (high P) relative to the estimated TER of a triploid *P. antipodarum* lineage of 270 C:P (Tibbets et al. 2010). We produced the high and low-P diets by manipulating the carbon: phosphorus (C:P) ratio of cultured *Scenedesmus obliquus* (green algae) following methods used in several earlier studies (e.g., Neiman et al. 2013b, Krist et al. 2014, Neiman and Krist 2016, Krist et al. 2017). We supplied 0.0004 g algae per snail three times a week following Neiman et al. (2013b). We measured individual snails weekly as described above. Individual snails were housed in a polystyrene cup filled with ∼250 mL of carbon-filtered tap water. We cleaned cups and changed water twice weekly and supplied each cup with a pinch of powdered chalk once per week to maintain shell growth. We checked weekly for dead snails and replaced dead or missing snails within the first week of the experiment with a similarly sized member of the same lineage.

We measured shell length of snails to assess whether specific growth rate (SGR) of juvenile *P. antipodarum* was affected by ploidy, P treatment, or their interaction. We calculated specific growth rate (SGR; day ^-1^) as ln(M_t_/M_o_)/t, where t is time, M_o_ is initial mass, and M_t_ is mass at 16 days. We calculated mass from shell length using the mass-length regression mass (mg) = 0.0199*L^2.35, where L = shell length (Hall et al. 2006). Because SGR is a measure of exponential growth, we measured SGR at the beginning of the experiment (from day 0 to 16) before growth rate slowed as snails approached reproductive maturity (Winterbourn 1970, Larkin et al. 2016).

### P content

Starting 9 weeks into the experiment, we began destructively sampling snails to measure P content once they exceeded 3.0 mm (triploids) or 3.33 mm (diploids); we applied a higher size threshold to diploids to ensure we avoided dissecting juvenile (unsexable) males. Because reproductive status can affect P content of female snails (Najev et al. 2026), we dissected all female snails to assess brooding status. We dissected 60 diploids and 64 triploids (N = 124 females), with 23 females harboring embryos. For brooding females, we combined the embryo tissue with the maternal tissue into a single sample when we measured P content. Prior to analyzing P content, we placed each sample on a glass fiber filter (Whatman, 24 mm) and dried the samples in a drying oven at 120°F for at least one week.

To prepare samples for analysis, we ashed all samples in a muffle furnace at 500°C for at least 2 hours and then digested the ashed sample in nitric acid in a Milestone GmbH ultraWAVE microwave digestion system. We used inductively coupled plasma mass spectrometry (ICP-MS) on an Agilent 7800 Quadrupole ICP-MS (Hachioji-shi, Tokyo, Japan) to measure the % P of each sample (P (mg)/ sample (mg)) hereafter “P content”. We ashed samples in a muffle furnace and conducted microwave digestions and mass spectrometry in the University of Iowa MATFab facility. After removing one missing and one mislabeled sample, we analyzed P content of 136 snails.

## Statistical Analysis

We used R software (4.4.2; R Core Team 2025) for all analyses. We ran general linear models using the “glm” function from the package “lme4” (Bates et al. 2015). We tested whether models met assumptions using “plotResiduals” function from the package “DHARMa” (Hartig 2018). We described outcomes of formal analyses following Wasserstein et al. (2019) and Muff et al. (2022) to avoid using an arbitrary probability threshold (e.g., 0.05).

### Specific Growth Rate

To evaluate whether individual growth rate differed as a function of ploidy, P limitation, or their interaction, we used a GLM with a gaussian distribution with an identity link, for specific growth rate (dependent variable), ploidy (diploid vs. triploid) and P treatment (low-vs. high-P diet) as fixed factors, and the interaction of ploidy by P treatment. We tested all assumptions with the DHARMA package, and all were met in our model. We removed all snails that died or were lost during the experiment or were not sexed. Altogether, we included 121 snails in analyses of growth rate, with 66 snails in the high-P treatment and 55 snails in the low-P treatment.

### P content

To evaluate whether snail P content differed in response to P limitation and/or ploidy, we used a GLM with a gaussian distribution with an identity link, with P content as the dependent variable, ploidy (diploid vs. triploid) and P treatment (low-vs. high-P diet) as fixed factors, and the interaction of ploidy by P treatment. Because *P. antipodarum* embryos are more P-rich than adult snails (Najev et al. 2026), we included a binary covariate of brooding vs. non-brooding. We also combined maternal and embryo P content (i.e., P for each brooding snail was the sum of mother P + embryo P) for this analysis. The model met all assumptions of a GLM. After removing all dead and missing snails from the analyses, we analyzed the P content of 124 snails: 68 snails in the high-P treatment and 56 snails in the low-P treatment.

## Results

### P content

Ploidy strongly influenced snail P content, with 14% higher P content in asexual triploids than sexual diploids (Figure 1, Table 1, S1). P content was not influenced by dietary P treatment, reproductive status, or the interaction between ploidy and P treatment (Table 1). Asexual triploids fed a high-P diet had the highest P content and sexual diploids fed a low-P diet had the lowest P content (Figure 1, Table 1).

**Figure 1.** P content (as %P) was higher in triploids than diploids but was not affected by dietary P treatment. Boxplots show the third (75th percentile) and first quartile (25th percentile) of the data, the horizontal line in the box is the median, and whiskers show the full range of the data. The violin shape depicts the distribution of the data, with the widest region depicting the most frequent values.

**Table 1.** GLM predicting P content by P treatment, Ploidy, Reproductive status (brooding or non-brooding) and the interaction between P treatment and ploidy (N = 124, 23 brooding snails and 101 non-brooding snails). For reproductive status, P content of brooding females includes their broods (embryos).

| Source | $X^2$ | df | $p$ |
| --- | --- | --- | --- |
| P treatment | 0.058 | 1 | 0.809 |
| Ploidy | 5.089 | 1 | 0.024 |
| Reproductive status | 0.138 | 1 | 0.711 |
| P treatment by Ploidy | 0.210 | 1 | 0.646 |

### Specific Growth Rate

Ploidy very strongly influenced growth rate (Figure 2, Table 2); asexual triploids grew 48% faster than sexual diploids (Table S2). We found no evidence that growth rate was influenced by P treatment nor by the interaction between ploidy and P treatment (Table 2). Asexual triploids fed a high-P diet had the highest growth rate and sexual diploids fed either diet had the lowest growth rate (Figure 2, Table S2).

**Figure 2.** Specific growth rate (day ^-1^) for the first 16 days was higher in triploids but not affected by dietary P treatment. Boxplots show the third (75th percentile) and first quartile (25th percentile) of the data, the horizontal line in the box is the median, and whiskers show the full range of the data. The violin shape depicts the distribution of the data, with the widest region depicting the most frequent values.

**Table 2.** GLM predicting specific growth rate (day ^-1^) as a function of P treatment, ploidy, and the interaction between P treatment and ploidy (N = 121).

| Source | $X^2$ | df | $p$ |
| --- | --- | --- | --- |
| P treatment | 0.005 | 1 | 0.947 |
| Ploidy | 14.126 | 1 | <0.001 |
| P treatment by Ploidy | 0.438 | 1 | 0.508 |

## Discussion

We examined whether *P. antipodarum* P content differed with ploidy level (Neiman et al. 2009), or with dietary P treatment (Tibbets et al. 2010, Neiman et al. 2013b) as expected and previously demonstrated. Comparisons of P content were our starting point because only if triploids possess higher P content than diploids would we expect greater sensitivity of triploid to limited dietary P availability. Our main goal was to provide a qualitative extension of previous work indicating that ploidy elevation is associated with higher sensitivity to P limitation of key life history traits in *Potamopyrgus antipodarum*. This earlier work showed that tetraploid asexual *P. antipodarum* are more severely affected by a P-limited diet than triploid asexual conspecifics (Neiman et al. 2013b) and that triploid asexuals grow faster than diploid sexuals when P is not limiting (Larkin et al. 2016, Gibson et al. 2017). However, what remained unclear is whether 1) growth of triploid asexuals is also more severely affected by P limitation than diploid sexuals, 2) whether triploids consistently grow faster than diploids when P is not limiting, and 3) whether the growth-rate advantage of asexual triploid *P. antipodarum* - if any - relative to sexual diploid conspecifics disappears when P is limiting. Answering these questions is also directly relevant to whether and to what extent differences in sensitivity to P limitation between the triploid asexuals and diploid sexuals could be relevant to the maintenance of sexual reproduction in natural populations of these snails.

### P content was highest in triploids but not affected by dietary P treatment

Because polyploid *P. antipodarum* have higher nuclear genome DNA content (Neiman et al. 2011, Neiman et al. 2025), more nucleic acids, and more P per unit mass than diploids (Neiman et al. 2009), we expected that the triploid snails would have higher P content than the diploids in our experiment, especially in the high-P treatment. We also expected P content to differ between diploids and triploids because tetraploid *P. antipodarum* had higher P content than triploids when P was adequate (Neiman et al. 2013b). Consistent with our predictions, triploids did have higher P content than diploids. But counter to our predictions, and in contrast to previous work and other studies in snails (e.g., Chaitanawisuti et al. 2010), we did not detect an effect of dietary P treatment on P content. Instead, the P content of triploids was similarly higher than diploids across both P treatments. One potential explanation for this result is that the increased allocation to P demanded by the 50% higher nucleic acid content of the asexual triploids (Neiman et al. 2009) obscures any more subtle effect of P diet on body composition when comparing between triploids and diploids.

### Growth rate of triploids is not more severely affected by P limitation than diploids

Experimental results also differed from expectations because, despite higher P content in triploids than diploids, growth of asexual triploids was not more severely affected by P limitation than sexual diploids in our experiment. Our result also contrasts with a previous study showing that tetraploid growth was more severely affected by P limitation than triploid growth (Neiman et al. 2013b). Together, these data indicate that ploidy and/or reproductive mode is a more important mediator of growth than P availability in *P. antipodarum*, suggesting that phosphorus availability in nature might not be a major contributor to the maintenance of ploidy polymorphism and sex.

Our results are broadly congruent with a substantial body of previous work with triploid *P. antipodarum* revealing substantial genetic and population-level variation of P limitation on individual growth rate. This variation is interesting and important in revealing a genetic, and potentially heritable, basis for sensitivity to P limitation but also necessarily reduces our likelihood of detecting effects of P limitation. For example, in Krist et al. (2016), only half (2 of 4) of the lineages grew more slowly on the P-limited than the P-sufficient diets. Similarly, despite a significant effect of dietary P treatment on growth of triploids, responses varied among lineages (Neiman et al. 2013b) and populations (Krist et al. 2017) from no response to a 50% reduction in growth rate on the P-limited diet relative to the P-sufficient diet. In another experiment with triploid lineages, we found no main effect of dietary P treatment on growth rate among 10 lineages (Krist et al. 2017) because some lineages grew more slowly on a P-limited diet while others did not. Major influences of genetic background on growth rate of triploid *P. antipodarum* have also been reported when P is not limiting (Larkin et al. 2016, Donne et al. 2022). Altogether, in past experiments, we have consistently found large effects of genetic background (lineage) on growth rate and response to P limitation, suggesting substantial variation in reaction norms in response to dietary P availability, which could mute or and even mask broad contrasts of the effects of P limitation across ploidy levels.

### Growth rate of triploids is higher than diploids independent of dietary P availability

Triploid snails possessed higher growth rates than diploids whether P availability was limiting or not. Our results are similar to earlier studies demonstrating higher growth rates in polypoid vs. diploid *P. antipodarum* (Larkin et al. 2016, Gibson et al. 2017) but extend the comparisons to include P availability. Higher growth rates in triploid relative to diploid conspecifics has also been reported in other molluscan taxa (e.g., bivalves, reviewed in Piferrer et al. 2009; *Crassostrea gigas* (Pacific oysters), Li and Li 2022) as well as in *Daphnia* (water flea, Jeyasingh et al. 2015), suggesting that increased growth rate could be a general positive consequence of increased ploidy in some aquatic invertebrates. Higher growth rates of triploids vs. diploids could be due to extra gene copies in polyploids permitting more rapid production of the mRNA, proteins, and ribosomes required for growth, though the extent to which polyploidy will *increase* organism-wide levels of these key molecules remains unclear (reviewed in Doyle and Coate 2019). Indirect support of this possibility include higher growth rates of polyploid *P. antipodarum* than diploid conspecifics (Larkin et al. 2016, Gibson et al. 2017), and higher rates of tissue regeneration reported in polyploid vs. diploid *P. antipodarum* (Krois et al. 2013). Nevertheless, the absence of a difference between triploid and tetraploid asexual snails (Krois et al. 2013, Larkin et al. 2016) indicates that ploidy elevation is not the whole story.

Asexual triploids may also grow faster than diploids because of strong and effective selection among clones representing the diverse assemblages of asexual *P. antipodarum* in many New Zealand lakes (Fox et al. 1996, Jokela et al. 2003, Paczesniak et al. 2013) selecting for high growth rates in asexual lineages. If genetic architecture for important traits like growth rate has an epistatic basis (e.g., Jakubowska and Korona 2012, Mackay 2014), then sexual *P. antipodarum* could be disadvantaged because recombination breaks up allelic combinations conferring rapid growth (Neiman and Linksvayer 2006). Because triploid asexual *P. antipodarum* are much more common and diverse than tetraploid asexual *P. antipodarum* (Neiman et al. 2011, Paczesniak et al. 2013), a larger role for interclonal selection is expected and could explain especially high performance of triploid relative to tetraploid conspecifics. While evidence that triploid outperform tetraploid *P. antipodarum* under lower but not higher P conditions (Neiman et al. 2013b) hints that ploidy plays at least a partial role in fitness-related aspects of the phenotype under some circumstances, rigorous characterization of the contributions of ploidy versus reproductive mode to *P. antipodarum* life history trait expression will ultimately require large experiments featuring a wide diversity of sexual diploid and asexual triploid and tetraploid *P. antipodarum*.

## Conclusions

Our study highlights the importance of ploidy in shaping life-history traits. Contrary to our predictions, we demonstrated performance advantages for asexual triploids over sexual diploids, even in P-limited conditions. This outcome is important in offsetting some or all disadvantages of polyploidy linked to P limitation. Superior growth performance of triploid asexual vs. diploid sexual *P. antipodarum* is also relevant to understanding the maintenance of sexual reproduction in this species. Our results provide yet another line of evidence that the “all else is equal” assumption between sexual vs. asexual females, expected to translate into a two-fold cost of males for sexual populations (Maynard-Smith 1978), is violated for *P. antipodarum* because asexual and sexual females are not equal in growth rate. Hence, our results make the maintenance of sex even more puzzling in this species because relative to asexual triploid populations, sexual populations face both the two-fold cost of males and inferior growth rates. Together, these data suggest that P availability will probably not play a major role in the maintenance of sexual diploid *P. antipodarum*.

## Acknowledgements

We thank Maricruz Mendoza, Vanessa Frimpong, Precious Pate, Chelsea Higgins, Humu Mohammed, Hannah Yerly, Bryan Guevara, Gwendolyn Gavin, Adi Castellano, Luca Angerer-Sueppel, and Waad Abdella for help with the experiment. We thank the Gillan lab at the University of Iowa for use of their muffle furnace. We also gratefully acknowledge the MATFab facility of the University of Iowa, specifically Linhan “Leo” Li and Micheal Sinwell.

## Data Availability Statement

Our data and code will be available as supplemental materials if our manuscript is accepted.

## Author contributions

● Briante Najev: Conceptualization, Methodology, Validation, Formal analysis, Investigation - Data collection, Resources, Data Curation, Writing - Original Draft, Writing - Review & Editing, Visualization, Supervision, Project administration, Funding acquisition
● Zachary Minthorn: Methodology, Validation, Formal analysis, Investigation - Data collection, Visualization, Supervision
● Seth Gordon: Investigation - Data collection, Supervision
● Josh Bliss: Methodology, Investigation - Data collection, Resources, Supervision
● Chase McInville: Methodology, Formal analysis, Investigation - Data collection, Resources, Supervision
● Vinny Chloros: Methodology, Investigation - Data collection, Resources, Data Curation, Visualization
● Waad Abdella: Methodology, Investigation - Data collection, Resources, Supervision
● Maurine Neiman: Conceptualization, Methodology, Validation, Formal analysis, Resources, Writing - Review & Editing, Visualization, Supervision, Project administration, Funding acquisition
● Amy C. Krist: Conceptualization, Methodology, Validation, Formal analysis, Resources, Writing - Review & Editing, Visualization, Supervision, Project administration, Funding acquisition

## References

1. Anneberg, J. T, and K. A. Segraves. 2020. Nutrient enrichment and neopolyploidy interact to increase lifetime fitness of *Arabidopsis thaliana*. Plant and Soil 456:439–453.

2. Anneberg, J. T, and K. A. Segraves. 2023. Neopolyploidy causes increased nutrient requirements and a shift in plant growth strategy in *Heuchera cylindrica*. Ecology 104:e4054.

3. Bates, D., M. Mächler, B. Bolker, and S. Walker. 2015. Fitting linear mixed-effects models using lme4. Journal of Statistical Software 67:1–48.

4. Cavalier -Smith, T. 2005. Economy, speed and size matter: evolutionary forces driving nuclear genome miniaturization and expansion. Annals of Botany 95:147–175.

5. Chaitanawisuti, N., T. Sungsirin, and S. Piyatiratitivorakul. 2010. Effects of dietary calcium and phosphorus supplementation on the growth performance of juvenile spotted babylon *Babylonia areolata* culture in a recirculating culture system. Aquaculture International 18:303–313.

6. Chen, Q., W. Zhu, L. Chang, M. Zhang, S. Wang, J. Liu, N. Lu, C. Li, F. Xie, B. Wang, and J. Jiang. 2025. Every gain comes with loss: ecological and physiological shifts associated with polyploidization in a pygmy frog. Molecular Biology and Evolution 42:msae258.

7. Cross, W. F., J. P. Benstead, P. C. Frost, and S. A. Thomas. 2005. Ecological stoichiometry in freshwater benthic systems: recent progress and perspectives. Freshwater Biology 50:1895–1912.

8. D’Souza, T. G., M. Storhas, and N. K. Michiels. 2005. The effect of ploidy level on fitness in parthenogenetic flatworms. Biological Journal of the Linnean Society 85:191–198.

9. Donne, C., K. Larkin, C. Adrian-Tucci, A. Good, C. Kephart, and M. Neiman. 2022. Life-history trait variation in native versus invasive asexual New Zealand mud snails. Oecologia 199:785–795.

10. Doyle, J. J., and J. E. Coate. 2019. Polyploidy, the nucleotype, and novelty: the impact of genome doubling on the biology of the cell. International Journal of Plant Sciences 180:1–52.

11. Du, E., C. Terrer, A. F. A. Pellegrini, A. Ahlström, C. J. van Lissa, X. Zhao, N. Xia, X. Wu, and R. B. Jackson. 2020. Global patterns of terrestrial nitrogen and phosphorus limitation. Nature Geoscience 13:221–226.

12. Elser, J. J., M. E. S. Bracken, E. E. Cleland, D. S. Gruner, W. S. Harpole, H. Hillebrand, J. T. Ngai, E. W. Seabloom, J. B. Shurin, and J. E. Smith. 2007. Global analysis of nitrogen and phosphorus limitation of primary producers in freshwater, marine and terrestrial ecosystems. Ecology Letters 10:1135–1142.

13. Elser, J. J., D. R. Dobberfuhl, N. A. MacKay, and J. H. Schampel. 1996. Organism size, life history, and N:P stoichiometry: toward a unified view of cellular and ecosystem processes. BioScience 46:674–684.

14. Fox, J. A., M. F. Dybdahl, J. Jokela, and C. M. Lively. 1996. Genetic structure of coexisting sexual and clonal subpopulations in freshwater snails (*Potamopygrus antipodarum*). Evolution 50:1541–1548.

15. Frost, P. C., J. P. Benstead, W. F. Cross, H. Hillebrand, J. H. Larson, M. A. Xenopoulos, and T. Yoshida. 2006. Threshold elemental ratios of carbon and phosphorus in aquatic consumers. Ecology Letters 9:774–779.

16. Geist, J. A., J. L. Mancuso, M. M. Morin, K. P. Bommarito, E. N. Bovee, D. Wendell, B. Burroughs, M. R. Luttenton, D. L. Strayer, and S. D. Tiegs. 2022. The New Zealand mud snail (*Potamopyrgus antipodarum*): autecology and management of a global invader. Biological Invasions 24:905–938.

17. Gerstein, A. C., H. Lim, J. Berman, and M. A. Hickman. 2017. Ploidy tug-of-war: evolutionary and genetic environments influence the rate of ploidy drive in a human fungal pathogen. Evolution 71:1025–1038.

18. Gibson, A. K., L. F. Delph, and C. M. Lively. 2017. The two-fold cost of sex: experimental evidence from a natural system. Evolution Letters 1:6–15.

19. Godfree, R. C., D. J. Marshall, A. G. Young, C. H. Miller, and S. Mathews. 2017. Empirical evidence of fixed and homeostatic patterns of polyploid advantage in a keystone grass exposed to drought and heat stress. R Soc Open Sci 4:170934.

20. Guignard, M. S., A. R. Leitch, C. Acquisti, C. Eizaguirre, J. J. Elser, D. O. Hessen, P. D. Jeyasingh, M. Neiman, A. E. Richardson, and P. S. Soltis. 2017. Impacts of nitrogen and phosphorus: from genomes to natural ecosystems and agriculture. Frontiers in Ecology and Evolution 5:70.

21. Guignard, M. S., R. A. Nichols, R. J. Knell, A. Macdonald, C.-A. Romila, M. Trimmer, I. J. Leitch, and A. R. Leitch. 2016. Genome size and ploidy influence angiosperm species’ biomass under nitrogen and phosphorus limitation. New Phytologist 210:1195–1206.

22. Hall Jr., R. O., M. F. Dybdahl, and M. C. VanderLoop. 2006. Extremely high secondary production of introduced snails in rivers. Ecological Applications 16:1121–1131.

23. Hartig, F. 2018. DHARMa: residual diagnostics for hierarchical (multi-level/mixed) regression models. R Package version 020.

24. Hessen, D. O., P. D. Jeyasingh, M. Neiman, and L. J. Weider. 2010. Genome streamlining and the elemental costs of growth. Trends in Ecology and Evolution 25:75–80.

25. Jakubowska, A., and R. Korona. 2012. Epistasis for growth rate and total metabolic flux in yeast. PLOS ONE 7:e33132.

26. Jeyasingh, P. D., P. Roy Chowdhury, M. W. Wojewodzic, D. Frisch, D. O. Hessen, and L. J. Weider. 2015. Phosphorus use and excretion varies with ploidy level in Daphnia. Journal of Plankton Research 37:1210–1217.

27. Jokela, J., and C. M. Lively. 1995. Parasites, Parasites, sex, and early reproduction in a mixed population of freshwater snails. Evolution 49:1268–1271.

28. Jokela, J., C. M. Lively, M. F. Dybdahl, and J. A. Fox. 2003. Genetic variation in sexual and clonal lineages of a freshwater snail. Biological Journal of the Linnean Society 79:165–181.

29. Krist, A. C., L. Bankers, K. Larkin, M. D. Larson, D. J. Greenwood, M. A. Dyck, and M. Neiman. 2017. Phosphorus availability in the source population influences response to dietary phosphorus quantity in a New Zealand freshwater snail. Oecologia 185:595–605.

30. Krist, A. C., A. D. Kay, K. Larkin, and M. Neiman. 2014. Response to phosphorus limitation varies among lake populations of the freshwater snail *Potamopyrgus antipodarum*. PLOS ONE 9:e85845.

31. Krist, A. C., A. D. Kay, E. Scherber, K. Larkin, B. J. Brown, D. Lu, D. T. Warren, R. Riedl, and M. Neiman. 2016. Evidence for extensive but variable nutrient limitation in New Zealand lakes. Evolutionary Ecology 30:973–990.

32. Krois, N. R., A. Cherukuri, N. Puttagunta, and M. Neiman. 2013. Higher rate of tissue regeneration in polyploid asexual versus diploid sexual freshwater snails. Biology Letters 9:20130422.

33. Larkin, K., C. Tucci, and M. Neiman. 2016. Effects of polyploidy and reproductive mode on life history trait expression. Ecology and Evolution 6:765–778.

34. Lewis, W. M. 1985. Nutrient scarcity as an evolutionary cause of haploidy. American Naturalist 125:692–701.

35. Li, Y., and Q. Li. 2022. The growth, survival and ploidy of diploid, triploid and tetraploid of the Pacific oyster (*Crassostrea gigas*) in larval and juvenile stages. Aquaculture 553:738083.

36. Lively, C. M., and J. Jokela. 2002. Temporal and spatial distributions of parasites and sex in a freshwater snail. Evolutionary Ecology Research 4:219–226.

37. Mackay, T. F. C. 2014. Epistasis and quantitative traits: using model organisms to study gene–gene interactions. Nature Reviews Genetics 15:22–33.

38. Maynard-Smith, J. 1978. The evolution of sex. Cambridge University Press, London, UK.

39. Morris, J. P., T. Baslan, D. E. Soltis, P. S. Soltis, and D. T. Fox. 2024. Integrating the study of polyploidy across organisms, tissues, and disease. Annual Review of Genetics 58:297–318.

40. Muff, S., E. B. Nilsen, R. B. O’Hara, and C. R. Nater. 2022. Rewriting results sections in the language of evidence. Trends in Ecology and Evolution 37:203–210.

41. Münzbergová, Z., and J. Skuhrovec. 2017. Contrasting effects of ploidy level on seed production in a diploid–tetraploid system. AoB Plants 9:plw077.

42. Najev, B., G. Gavin, C. Craven, A. Escandón, P. Pate, A. C. Krist, and M. Neiman. 2026. Phosphorus availability affects multiple metrics of female reproductive investment in a freshwater snail. Royal Society Open Science 13:251895.

43. Neiman, M., M. J. Beaton, D. O. Hessen, P. D. Jeyasingh, and L. J. Weider. 2017. Endopolyploidy as a potential driver of animal ecology and evolution. Biological Reviews 92:234–247.

44. Neiman, M., A. D. Kay, and A. C. Krist. 2013a. Can resource costs of polyploidy provide an advantage to sex? Heredity 110:152–159.

45. Neiman, M., A. D. Kay, and A. C. Krist. 2013b. Sensitvity to phosphorus limitation increases with ploidy level in a New Zealand snail. Evolution 67:1511–1517.

46. Neiman, M., and A. Krist. 2016. Sensitivity to dietary phosphorus limitation in native vs. invasive lineages of a New Zealand freshwater snail. Ecological Applications 26:2218–2224.

47. Neiman, M., K. Larkin, A. R. Thompson, and P. Wilton. 2012. Male offspring production by asexual *Potamopyrgus antipodarum*, a New Zealand snail. Heredity 109:57–62.

48. Neiman, M., and T. A. Linksvayer. 2006. The conversion of variance and the evolutionary potential of restricted recombination. Heredity 96:111–121.

49. Neiman, M., D. Paczesniak, D. M. Soper, A. T. Baldwin, and G. Hehman. 2011. Wide variation in ploidy level and genome size in a New Zealand freshwater snail with coexisting sexual and asexual lineages. Evolution 65:3202–3216.

50. Neiman, M., M. Pichler, M. Haase, and D. K. Lamatsch. 2025. Characterization of nuclear genome size and variation in a freshwater snail model system featuring a recent whole-genome duplication. Royal Society Open Science 12:250171.

51. Neiman, M., and T. Schwander. 2011. Using parthenogenetic lineages to identify advantages of sex. Evolutionary Biology 38:115–123.

52. Neiman, M., K. M. Theisen, M. E. Mayry, and A. D. Kay. 2009. Can phosphorus limitation contribute to the maintenance of sex? A test of a key assumption. Journal of Evolutionary Biology 22:1359–1363.

53. Otto, S. P., and J. Whitton. 2000. Polyploid incidence and evolution. Annual Review of Genetics 34:401–437.

54. Paczesniak, D., J. Jokela, K. Larkin, and M. Neiman. 2013. Discordance between nuclear and mitochondrial genomes in sexual and asexual lineages of the freshwater snail *Potamopyrgus antipodarum*. Molecular Ecology 22:4695–4710.

55. Phillips, N. R., and D. M. Lambert. 1989. Genetics of *Potamopyrgus antipodarum* (Gastropoda: Prosobranchia): evidence for reproductive modes. New Zealand Journal of Zoology 16:435–445.

56. Piferrer, F., A. Beaumont, J.-C. Falguière, M. Flajšhans, P. Haffray, and L. Colombo. 2009. Polyploid fish and shellfish: production, biology and applications to aquaculture for performance improvement and genetic containment. Aquaculture 293:125–156.

57. R Core Team. 2025. R: a language and environment for statistical computing. R Foundation for Statistical Computing, Vienna, Austria.

58. Riaz, S., Y. Jiang, M. Xiao, D. You, A. Klepacz-Smółka, F. Rasul, and M. Daroch. 2022. Generation of miniploid cells and improved natural transformation procedure for a model cyanobacterium *Synechococcus elongatus* PCC 7942. Frontiers in Microbiology 13:959043.

59. Saccucci, M. J., R. D. Denton, M. L. Holding, and H. L. Gibbs. 2016. Polyploid unisexual salamanders have higher tissue regeneration rates than diploid sexual relatives. Journal of Zoology 300:77–81.

60. Tibbets, T. M., A. C. Krist, R. O. Hall, and L. A. Riley. 2010. Phosphorus-mediated changes in life history traits of the invasive New Zealand mudsnail (*Potamopyrgus antipodarum*). Oecologia 163:549–559.

61. Turcotte, M. M., N. Kaufmann, K. L. Wagner, T. A. Zallek, and T.-L. Ashman. 2024. Neopolyploidy increases stress tolerance and reduces fitness plasticity across multiple urban pollutants: support for the “general-purpose” genotype hypothesis. Evolution Letters 8:416–426.

62. van de Pol, I. L. E., A. Hermaniuk, and W. C. E. P. Verberk. 2021. Interacting effects of cell size and temperature on gene expression, growth, development and swimming performance in larval zebrafish. Frontiers in Physiology 12:738804.

63. Veselý, P., P. Bureš, and P. Šmarda. 2013. Nutrient reserves may allow for genome size increase: evidence from comparison of geophytes and their sister non-geophytic relatives. Annals of Botany 112:1193–1200.

64. Wallace, C. 1992. Parthenogenesis, sex, and chromosomes in *Potamopyrgus*. Journal of Molluscan Studies 58:93–107.

65. Wang, X., J. A. Morton, J. Pellicer, I. J. Leitch, and A. R. Leitch. 2021. Genome downsizing after polyploidy: mechanisms, rates and selection pressures. Plant Journal 107:1003–1015.

66. Wasserstein, R. L., A. L. Schirm, and N. A. Lazar. 2019. Moving to a world beyond “p < 0.05”. The American Statistician 73:1–19.

67. Wendel, J. F. 2015. The wondrous cycles of polyploidy in plants. American Journal of Botany 102:1753–1756.

68. Winterbourn, M. J. 1970. Population studies on the New Zealand freshwater gastropod, *Potamopyrgus antipodarum* (Gray). Journal of Molluscan Studies 39:139–149.

