## Supplementary material for "Triploid asexual freshwater snails grow faster than sexual diploid conspecifics regardless of dietary phosphorus availability": Figures

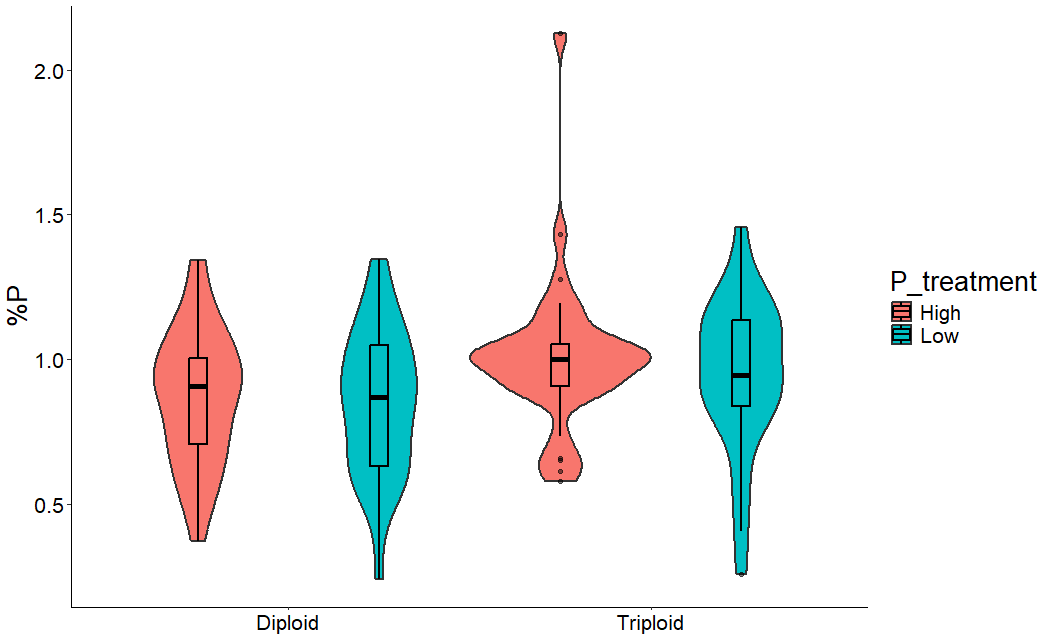
Figure 1.

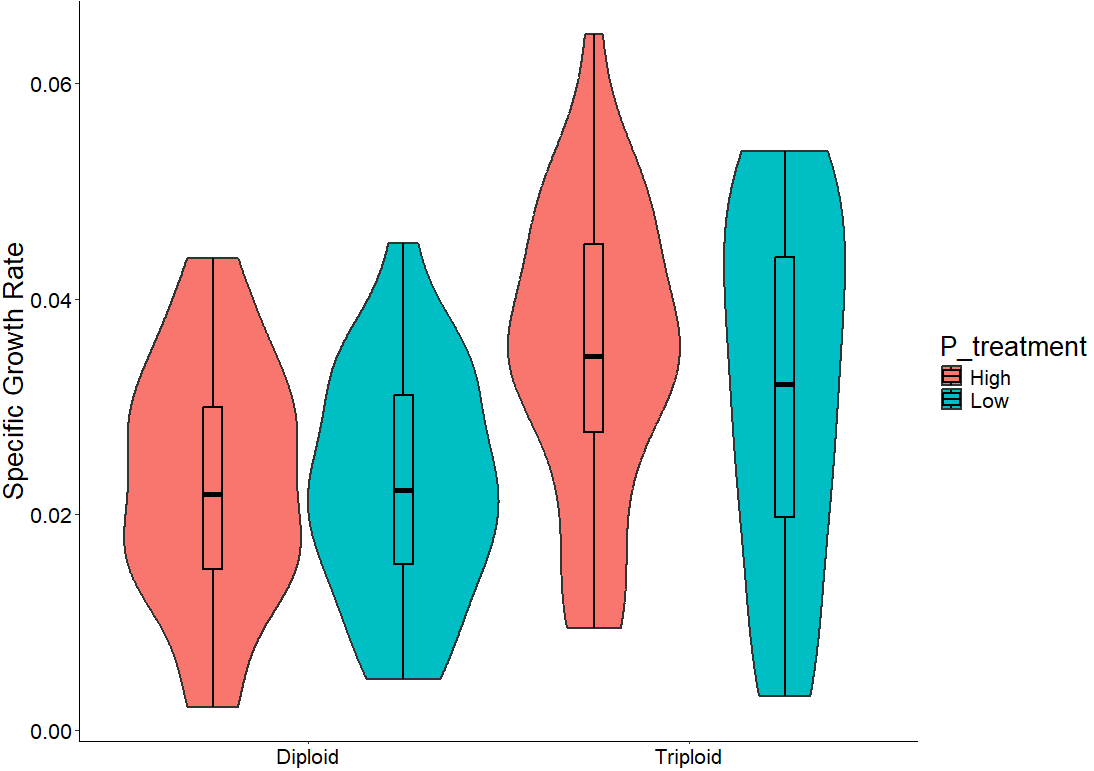

Figure 2.
