## Supplemental Tables for "Triploid asexual freshwater snails grow faster than sexual diploid conspecifics regardless of dietary phosphorus availability"

Table S1: Descriptive statistics for P content by ploidy and P treatment (N = 124).

| P treatment | Ploidy | Mean | Median | Standard deviation | Minimum value | Maximum value |
| --- | --- | --- | --- | --- | --- | --- |
| High | Diploid | 0.868 | 0.905 | 0.232 | 0.370 | 1.343 |
| High | Triploid | 1.007 | 1.001 | 0.263 | 0.579 | 2.126 |
| Low | Diploid | 0.850 | 0.868 | 0.252 | 0.238 | 1.346 |
| Low | Triploid | 0.949 | 0.946 | 0.265 | 0.256 | 1.457 |

Table S2: Descriptive statistics for specific growth rate (day ^-1^) by ploidy and P treatment (N = 121).

| P treatment | Ploidy | Mean | Median | Standard deviation | Minimum value | Maximum value |
| --- | --- | --- | --- | --- | --- | --- |
| High | Diploid | 0.023 | 0.022 | 0.011 | 0.002 | 0.044 |
| High | Triploid | 0.035 | 0.035 | 0.014 | 0.009 | 0.065 |
| Low | Diploid | 0.023 | 0.022 | 0.010 | 0.005 | 0.045 |
| Low | Triploid | 0.032 | 0.032 | 0.016 | 0.003 | 0.054 |
